# Intestinal Dysbiosis in Necrotic Enteritis: Dissecting the Roles of *Eimeria* and *Clostridium perfringens*

**DOI:** 10.64898/2026.05.27.728132

**Authors:** Jing Liu, Jiaqing Guo, Melanie A. Whitmore, Isabel Tobin, Dohyung M. Kim, Guolong Zhang

## Abstract

Necrotic enteritis (NE), caused by *Clostridium perfringens,* is a major enteric disease in poultry with substantial economic impact. NE is frequently triggered by co-infection with *Eimeria spp*., yet the relative contributions of *Eimeria* and *C. perfringens* to NE-induced dysbiosis and disease progression remain poorly defined. To address this, Cobb broiler chickens were challenged with *Eimeria maxima*, *C. perfringens*, or both, and ileal and cecal microbiota were analyzed using 16S rRNA gene sequencing and shotgun metagenomics. Temporal dynamics of intestinal microbiota shifts were further assessed at multiple time points post-infection. Our findings demonstrate that NE-associated dysbiosis is characterized by enrichment of pathobionts including *C. perfringens*, *Escherichia*, and *Enterococcus cecorum*, along with differential abundance of certain lactic acid-and short chain fatty acid-producing bacteria. Dysbiosis and disease progression were driven by synergistic interactions between *E. maxima* and *C. perfringens*, with *E. maxima* exerting a dominant influence. Notably, *E. maxima* alone promoted expansion of commensal *C. perfringens* or closely-related bacteria, even without prior exposure. Metagenomic analysis of the cecal microbiota further revealed a functional shift favoring utilization of host-derived glycans and simple carbohydrates over dietary fibers, in response to *E. maxima* and NE challenges. This transition coincided with *E. maxima*-induced epithelial damage, increased mucogenesis, and nutrient malabsorption. NE-associated dysbiosis emerged four days post-*E. maxima* infection and peaked 2–3 days following *C. perfringens* challenge. These findings suggest that *Eimeria* infection creates a permissive intestinal environment for *C. perfringens* colonization and proliferation, underscoring its pivotal role in NE pathogenesis.

**IMPORTANCE:** NE is a major health and economic burden in poultry production, primarily driven by *C. perfringens* and potentiated by *Eimeria* infection. This study provides critical mechanistic insights into the distinct and synergistic roles of *E. maxima* and *C. perfringens* in NE disease progression. We show that *Eimeria* plays a dominant role in driving NE-associated dysbiosis and disease progression by inducing epithelial damage, inflammation, and nutrient malabsorption, which facilitate *C. perfringens* colonization, proliferation, and toxin production. Comprehensive characterizations of both structural and functional microbiome shifts revealed that NE-associated dysbiosis is marked by enrichment of facultative pathobionts that favors utilization of host-derived mucins and simple dietary carbohydrates, alongside depletion of strictly anaerobic, fiber-fermenting, SCFA-producing bacteria. These microbial shifts reflect disease progression and offer potential biomarkers for early diagnosis and therapeutic intervention. Our findings lay a foundation for microbiota-based diagnostics and interventions to mitigate NE and other enteric diseases.

## INTRODUCTION

Necrotic enteritis (NE) is an important poultry disease, responsible for an estimated global economic loss of approximately $6 billion annually (1). NE can be classified into two forms: clinical and subclinical. Clinical NE is characterized by severe intestinal lesions, diarrhea, microbiota shifts, and high mortality rates that can reach up to 50% (2). In contrast, subclinical NE often goes undetected but leads to decreased weight gain and feed efficiency. Due to its high prevalence, subclinical NE exerts a substantially greater economic impact on animal productivity and profitability than the clinical form. The etiological agent of NE is *Clostridium perfringens* (CP), whose proliferation and toxin production result in inflammation and damage to the small intestinal mucosa (2). However, CP infection alone typically results in minimal pathology in healthy chickens. Predisposing factors such as coccidiosis, stress, immunosuppression, and dietary and environmental conditions are critical for the development of NE (3).

*Eimeria spp.*, the causative agents of coccidiosis, are recognized as a major predisposing factor for NE in commercial poultry production (2, 3). *Eimeria* infection causes epithelial damage, exposing extracellular matrix components such as collagen, which serve as binding sites for CP and facilitate the production of virulence factors like NetB toxin (4). Additionally, *Eimeria* infection stimulates mucus production, creating a nutrient-rich environment that supports the proliferation of mucolytic bacteria, including *C. perfringens* (5). Consequently, *Eimeria maxima* (EM), which targets the small intestine, is commonly used in experimental NE models to predispose chickens to CP infection (2, 3).

The gastrointestinal tract (GIT) of chickens harbors a complex and diverse microbiota that plays vital roles in nutrient digestion and absorption, immune development, and colonization resistance against pathogens. NE is associated with extensive alternations in the intestinal microbiota, often marked by a reduction in short-chain fatty acid (SCFA)-producing bacteria and an enrichment of pathobionts such as *Escherichia* and *Enterococcus* (6-8). Although co-infection with CP and EM has been shown to disrupt microbial diversity and composition, the individual contributions of each pathogen to NE-associated dysbiosis remain poorly defined (9-12).

This study aims to dissect the specific roles of CP and EM in driving intestinal dysbiosis and disease progression in NE. By characterizing the structural and predicted functional changes in the ileal and cecal microbiota following single and combined infections, we seek to elucidate the microbial dynamics underlying NE disease progression. These insights will inform the development of microbiome-based diagnostic and therapeutic strategies for NE and other enteric diseases in poultry.

## RESULTS

### Differential response of chickens to CP, EM, and NE infections

To evaluate relative contributions of CP and EM to NE-induced dysbiosis, we orally challenged Cobb broiler chickens with CP, EM, or both pathogens to induce NE (**Fig. 1A**). CP infection alone resulted in no mortality, while EM challenge led to a survival rate of approximately 93%. In contrast, co-infection with CP and EM significantly reduced survival to 26% (**Fig. 1B**). Among the 39 surviving co-infected chickens, 18 exhibited mild intestinal lesions (score 1), 17 showed severe necrotic lesions along the jejunum and ileum (score 6), and four had moderate lesions (scores 2–3) based on a 6-point scoring system (13).

**Fig. 1.**
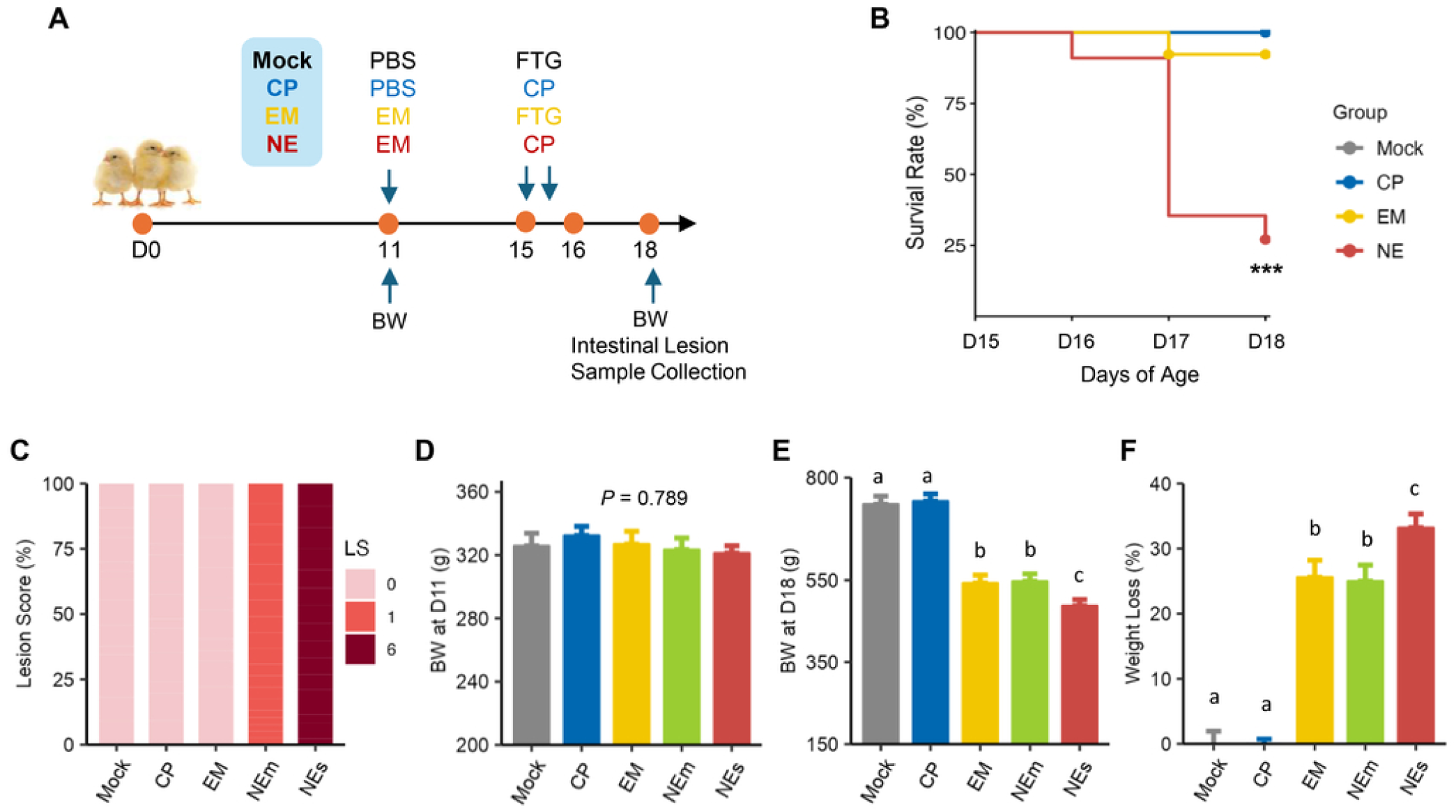
Differential response of chickens to infections. (**A**) Experimental design of Trial 1. A total of 195 day-of-hatch male Cobb broilers were orally inoculated with *E. maxima* (EM; *n* = 15), *C. perfringens* (CP; *n* = 15, administered twice), or both pathogens to induce necrotic enteritis (NE; *n* = 150) on days 11 and 15, respectively. A fourth group (*n* = 15) were mock-infected with PBS and fluid thioglycollate broth (FTG) on respective days. Animal survival was recorded twice daily till day 18. Individual body weights (BW) were recorded on days 11 and 18. All surviving birds were humanely euthanized on day 18, and small intestinal lesions were scored on a scale from 0 to 6. (**B**) Survival rates from day 15 to day 18. Statistical significance was determined using the log-rank test. \*\*\**P* < 0.001 vs. mock-infected group. (**C**) Distribution (%) of small intestinal lesion scores (LS) among surviving birds on day 18. Surviving animals in the NE group were further stratified into two subgroups: NE with mild, score-1 lesions (NEm; *n* = 18) and NE with severe, score-6 lesions (NEs; *n* = 17). (**D–F**) Average BW on day 11, BW on day 18, and weight loss (%) between day 11 and 18 in surviving birds. Data are presented as mean ± SEM. Groups not sharing a common superscript letter differ significantly (*P* < 0.05) based on one-way ANOVA followed by post-hoc Tukey test.

Given marked microbiome differences between mild and severe NE cases (7), we stratified co-infected birds into mildly affected (NEm, score 1) and severely affected (NEs, score 6) groups for downstream microbiome analyses. The four birds with intermediate lesion scores were excluded due to limited sample size. No NE-characteristic lesions were observed in the mock, CP, or EM groups, all of which received a lesion score of 0 (**Fig. 1C**). Although all groups began with comparable body weights (**Fig. 1D**), significant growth retardation was observed in the EM, NEm, and NEs groups by the end of the trial, but not in the CP group (**Fig. 1E, 1F**).

### Alterations in ileal microbiota composition in response to CP, EM, and NE infections

To dissect relative contributions of CP and EM to NE-induced dysbiosis, we performed 16S rRNA gene sequencing on bacterial DNA isolated from ileal and cecal digesta. Following quality control, 58,543,086 high-quality sequencing reads were obtained with an average of 107,814 ± 862 sequences per sample. After filtering out low-prevalence amplicon sequence variants (ASVs) present in < 5% of samples, 405 and 443 ASVs were retained from the ileal and cecal samples, respectively. In the ileum, microbiota richness, evenness, and diversity, as measured by Observed ASVs (**Fig. 2A**), Pielou’s Evenness (**Fig. 2B**), and Shannon Index (**Fig. 2C**), respectively, were significantly reduced in the EM, NEm, and NEs groups, but not in the CP group, relative to the mock group. Weighted UniFrac analysis revealed significant differences (*P* < 0.05) in bacterial community structure across groups (**Fig. 2D**). Pairwise comparisons further highlighted significant differences in weighted UniFrac distance for nearly each pair, except between the mock and CP groups (**Table S1A**), suggesting that CP alone had minimum impact, while EM and NE induced more pronounced changes.

**Fig. 2.**
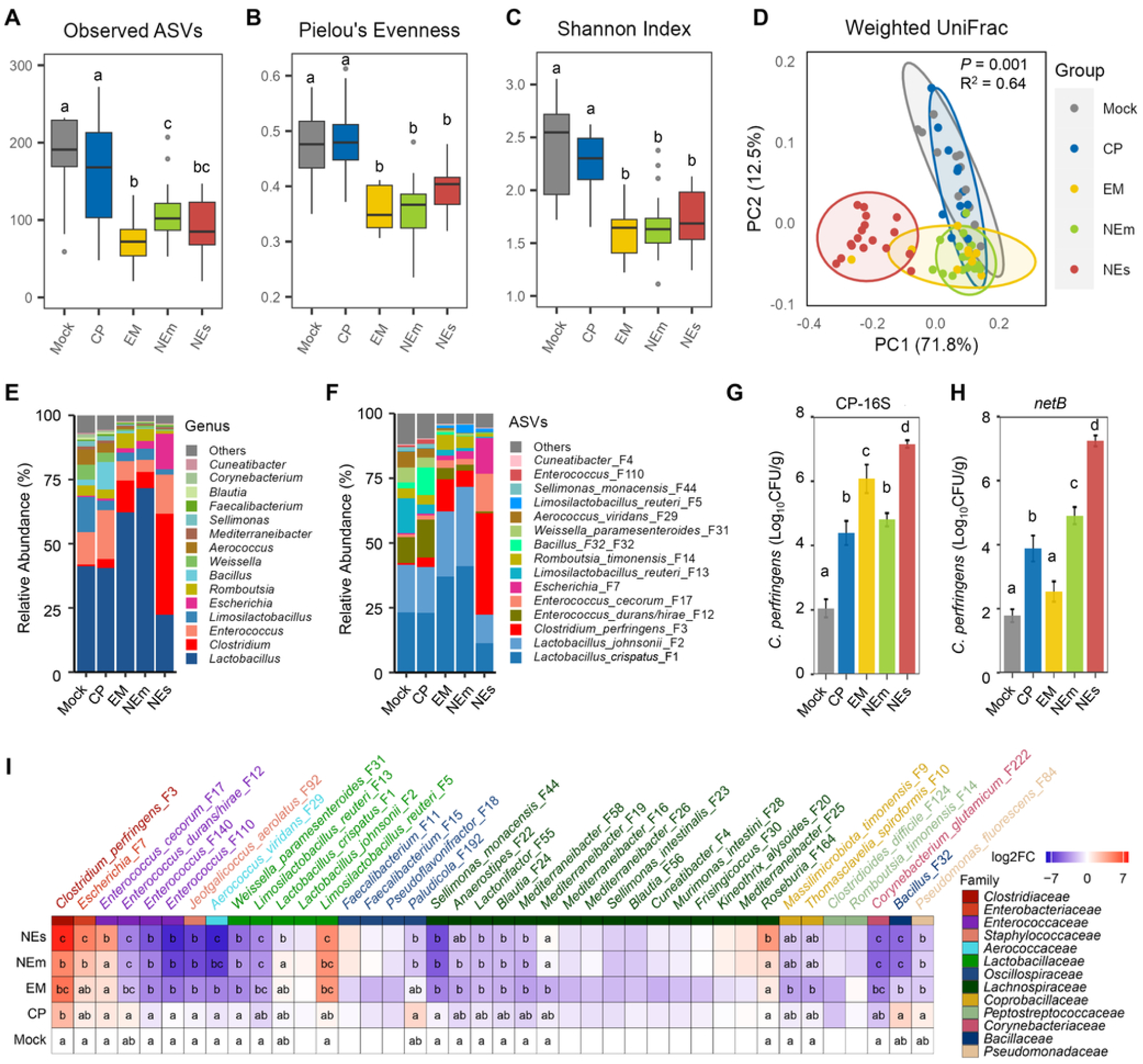
Alterations in ileal microbiota composition in response to infections. In Trial 1, day-of-hatch male Cobb broilers were mock-infected (*n* = 15) or orally challenged with *E. maxima* (EM; *n* = 15), *C. perfringens* (CP; *n* = 15), or both pathogens to induce necrotic enteritis (NE). Surviving animals in the NE group were further stratified into two subgroups: NE with mild, score-1 lesions (NEm; *n* = 18) and NE with severe, score-6 lesions (NEs; *n* = 17). Ileal digesta samples were collected from surviving animals on day 18 for DNA isolation and 16S rRNA gene sequencing. **(A–C)** Alpha diversity metrics including observed amplicon sequence variants (ASVs), Pielou’s Evenness, and Shannon Index, visualized using box-and-whisker plots. Statistical significance was assessed using the Kruskal–Wallis test followed by post-hoc Dunn’s test. Groups not sharing a common superscript letter differ significantly (*P* < 0.05). (**D**) Principal coordinates analysis (PCoA) plots of weighted UniFrac distances. Significance was determined using PERMANOVA using 999 permutations. (**E–F**) Relative abundances (%) of the top 15 genera and top 15 ASVs in the ileal microbiota. (**G–H**) Quantification of *C. perfringens* titers in ileal digesta using bacterium-specific 16S rRNA gene primers and *netB*-specific primers. Data are presented as mean ± SEM. Statistical comparisons were made using one-way ANOVA followed by post-hoc Tukey test. Groups not sharing a common superscript letter differ significantly (*P* < 0.05). (**I**) Heatmap depicting differential enrichment of the top 40 ASVs in the ileum. Data are shown as log_2_-transformed fold changes relative to mock-infected controls. Statistical significance was determined using ANCOM-BC2 analysis. Groups not sharing a common superscript letter within a column differ significantly (*P* < 0.05).

Taxonomically, *Lactobacillus* was the dominant genus in the ileum, comprising 33.9–71.2% of the total bacterial population (**Fig. 2E)**. However, in NEs chickens, *Clostridium* became predominant, accounting for 39.5% of the community. At the ASV level, *Lactobacillus crispatus* (F1) and *Lactobacillus johnsonii* (F2) were the most abundant in the mock, CP, EM, and NEm groups, while major pathobionts including *C. perfringens* (F3), *Escherichia* (F7), and *Enterococcus cecorum* (F17) dominated in the NEs group (**Fig. 2F**). *Escherichia* (F7) and *E. cecorum* (F17) experienced progressive enrichment correlating with NE severity. *C. perfringens* (F3) was also markedly enriched in CP-infected chickens, with a progressive rise in relative abundance from 0.2% in the mock group to 1.7%, 4.6%, and 39.5% in the CP, NEm, and Nes chickens, respectively. To our surprise, *C. perfringens* significantly increased to 5.6% in EM-challenged chickens, despite no direct prior exposure to CP. To verify elevated *C. perfringens* was derived from a commensal source rather than cross-contamination, quantitative PCR (qPCR) with primers specific for the *C. perfringens* 16S rRNA gene detected a significant increase in *C. perfringens* in all groups, reaching 1.2 × 10^6^ CFU/g in the ileum of EM-challenged chickens (**Fig. 2G**), while *netB*-specific primers only detected approximately 347 CFU/g of *C. perfringens*, comparable the mock group (**Fig. 2H**). These results suggested that enriched *C. perfringens* in EM chickens is likely a commensal, *netB*-negative *C. perfringens* strain or a closely related *Clostridium* species.

Among the 40 most abundant ileal bacterial ASVs, CP infection caused mostly subtle changes compared to EM and NE infections. Notably, EM-induced changes closely resembled those observed in NE, particularly in the NEm group (**Fig. 2I**). Approximately two-thirds of the bacteria were differentially regulated by EM and NE infections, with CP infection mostly exerting a negligible impact. Within the dominant lactic acid bacteria (LAB), *L. crispatus* (F1) and *L. johnsonii* (F2) were slightly enriched in EM and NEm chickens but reduced in NEs chickens. Interestingly, two strains of *Limosilactobacillus reuteri* displayed opposing trends: F5 significantly increased, while F13 significantly decreased across EM, NEm, and NEs groups. Additionally, several minor LAB species showed differential enrichment patterns. *Weissella paramesenteroides* (F31) and *Aerococcus viridans* (F29) were significantly diminished in response to EM and NE infections, while four *Enterococcus* ASVs showed a polarized response. *E. cecorum* (F17) was elevated, but three other *Enterococcus* members (F12, F110, and F140) were suppressed. Although representing minor constituents of the ileal microbiota, SCFA-producing bacteria belonging to the families *Oscillospiraceae, Lachnospiraceae*, and *Peptostreptococcaceae* were largely reduced by EM or NE infections.

### Alterations in cecal microbiota composition in response to CP, EM, and NE infections

Compared to the ileal microbiota, the cecal microbiota exhibited relatively modest alterations in response to CP, EM, and NE infections. The α-diversity metrics, including species richness (**Fig. 3A**), evenness (**Fig. 3B**), and Shannon Index (**Fig. 3C**), showed no significant differences among groups (*P* > 0.05). However, β-diversity analysis using weighted UniFrac distances revealed significant shifts across groups (**Fig. 3D**), with most pairwise comparisons showing statistical significance (*P* < 0.05; **Table S1B**).

**Fig. 3.**
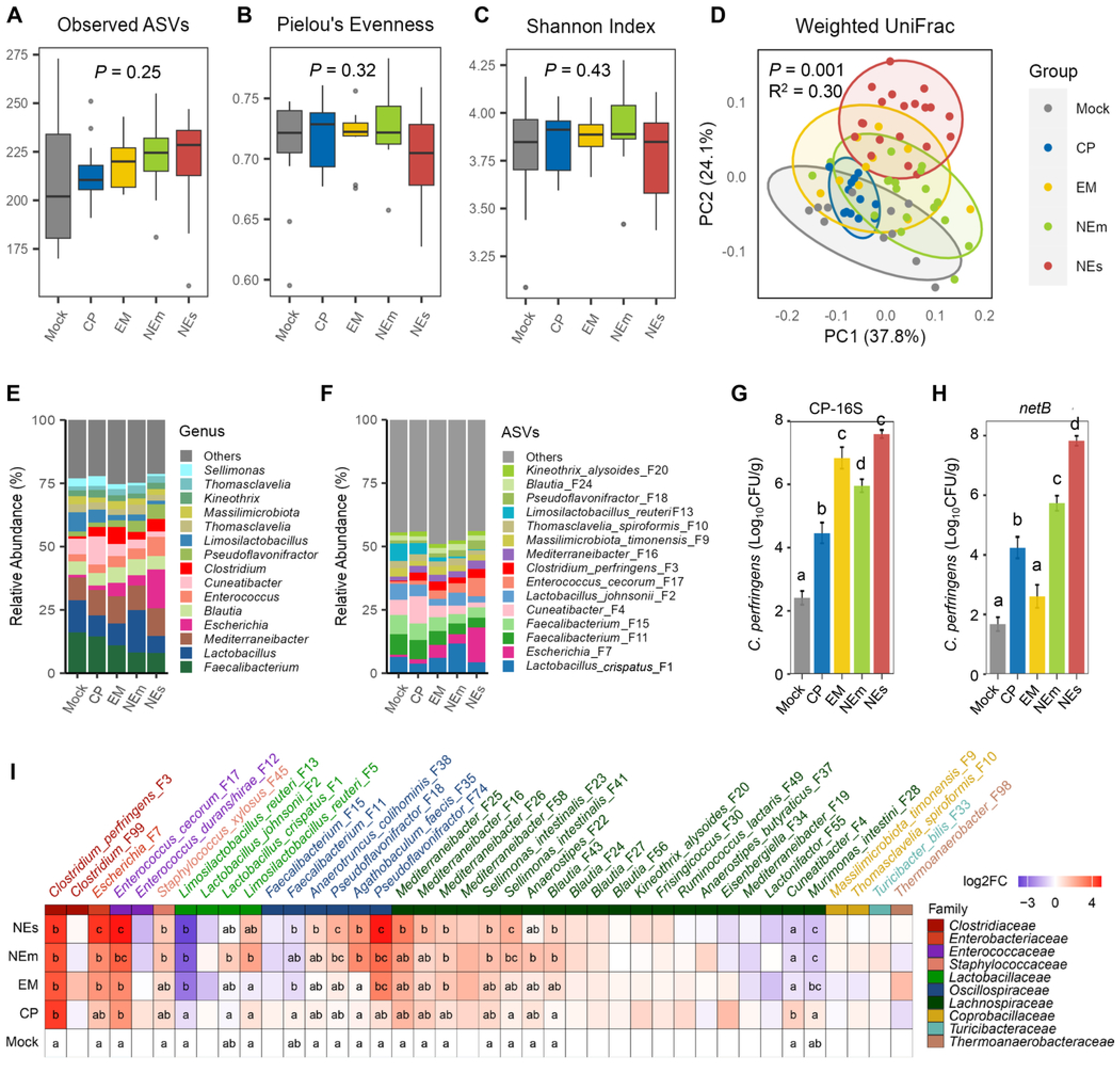
Alterations in cecal microbiota composition in response to infections. In Trial 1, day-of-hatch male Cobb broilers were mock-infected (*n* = 15) or orally challenged with *E. maxima* (EM; *n* = 15), *C. perfringens* (CP; *n* = 15), or both pathogens to induce necrotic enteritis (NE). Surviving animals in the NE group were further stratified into two subgroups: NE with mild, score-1 lesions (NEm; *n* = 18) and NE with severe, score-6 lesions (NEs; *n* = 17). Cecal digesta samples were collected from surviving animals on day 18 for DNA isolation and 16S rRNA gene sequencing. **(A–C)** Alpha diversity metrics including observed amplicon sequence variants (ASVs), Pielou’s Evenness, and Shannon Index, visualized using box-and-whisker plots. Statistical significance was assessed using the Kruskal–Wallis test followed by post-hoc Dunn’s test. Groups not sharing a common superscript letter differ significantly (*P* < 0.05). (**D**) Principal coordinates analysis (PCoA) plots of weighted UniFrac distances. Significance was determined using PERMANOVA using 999 permutations. (**E–F**) Relative abundances (%) of the top 15 genera and top 15 ASVs in the cecal microbiota. (**G–H**) Quantification of *C. perfringens* titers in cecal digesta using bacterium-specific 16S rRNA gene primers and *netB*-specific primers. Data are presented as mean ± SEM. Statistical comparisons were made using one-way ANOVA followed by post-hoc Tukey test. Groups not sharing a common superscript letter differ significantly (*P* < 0.05). (**I**) Heatmap depicting differential enrichment of the top 40 ASVs in the cecum. Data are shown as log_2_-transformed fold changes relative to mock-infected controls. Statistical significance was determined using ANCOM-BC2 analysis. Groups not sharing a common superscript letter within a column differ significantly (*P* < 0.05).

The 15 most abundant genera accounted for 59.1–72.6% of the total cecal bacterial population, with *Faecalibacterium*, *Lactobacillus*, *Mediterraneibacter*, and *Escherichia* being the most prevalent (**Fig. 3E**). The top 15 bacterial ASVs represented 36.4–50.7% of the total cecal bacteria, with *L. crispatus* (F1), *Escherichia* (F7) and two *Faecalibacterium* ASVs (F11 and F15) being the most prevalent (**Fig. 3F**). Similar to our observations in the ileal microbiota, *C. perfringens* was markedly enriched in the cecum of EM-challenged chickens. The qPCR analysis of the 16S rRNA and *netB* genes of *C. perfringens* again confirmed that increased *C. perfringens* in EM*-*infected chickens was primarily due to enrichment of commensal, *netB*-negative *C. perfringens* or closely related species (**Fig. 3G, 3H**).

Among the 40 most abundant cecal bacteria, EM challenge induced more pronounced shifts than CP (**Fig. 3I**). In addition to *C. perfringens*, two other major pathobionts—*Escherichia* (F7) and *E. cecorum* (F17)—were progressively enriched in EM, NEm, and NEs groups. In contrast, a different *Enterococcus* species (F12) was progressively reduced in NE chickens with increasing disease severity. Within LAB, *L. crispatus* (F1) was slightly enriched in NEm chickens, while *L. johnsonii* (F2) was reduced in EM and NEs chickens. Similar to the observations in the ileum, two *L. reuteri* ASVs displayed divergent responses, with F5 increasing in NEm and NEs chickens and F13 decreasing in CP, EM, NEm, and NEs groups. Two dominant SCFA-producing *Faecalibacterium* (F11 and F15) was reduced in response to EM and NE (**Fig. 3I**). In contrast, several other *Oscillospiraceae* members (F18, F35, F38, and F74) were enriched particularly in NEs chickens. Multiple SCFA-producing *Lachnospiraceae* members such as *Cuneatibacter* (F4) and *Murimonas intestini* (F28) progressively declined in EM- and NE-infected chickens, while others such as *Sellimonas intestinalis* (F23 and F41), *Blautia* (F43), and *Mediterraneibacter* (F16, F25, and F26) were enriched.

### Functional shifts in the cecal microbiome in response to CP, EM, and NE infections

To investigate functional alterations in the cecal microbiota in response to infections with CP, EM, or NE, we performed shotgun metagenomic sequencing on six cecal microbial DNA samples per treatment group. After filtering out host-derived reads, a total of 798 Gb of high-quality paired-end sequences were obtained, averaging 13.3 Gb per sample. Contigs from both individual and co-assembled metagenomes were binned, yielding 3,448 bins, of which 1,959 were of high quality, with estimated ≥ 80% completeness and ≤ 5% contamination. These high-quality bins were subsequently dereplicated at a 99% average nucleotide identity (ANI) threshold, resulting in 200 non-redundant metagenome-assembled genomes (MAGs). From these MAGs, we constructed a comprehensive non-redundant gene catalog comprising 849,390 unique genes, with 87.4% being classified into eggNOG clusters, 80% as Clusters of Orthologous Groups of proteins (COGs), 27.6 % to KEGG pathways, and 1.2% as carbohydrate-active enzymes (CAZymes).

To investigate potential involvement of metabolic pathways, we mapped bacterial genes to the KEGG database, identifying 477 pathways, of which 105 exhibited significant differential abundance. Among the top 12 pathways, those involved in transcriptional regulation, two-component systems, LPS biosynthesis, *E. coli* biofilm formation, bacterial toxin production, and amino acid metabolism were significantly enriched in both EM and NEs chickens (**Fig. 4A**). For example, multiple *C. perfringens* toxin genes such as *Plc/Cpa, PfoA, CloSI*, and *ColA*, as well as two major genes regulating the *C. perfringens* toxin production—*agrD* and *agrB*—were significantly enriched in EM, NEm, and/or NEs groups (**Fig. 4B, 4C**). Additionally, many genes involved in *E. coli* biofilm formation (**Fig. 4D**) and LPS biosynthesis (**Fig. 4E**) showed a significant increase in EM and NEs chickens (**Fig. 4B**). These results are consistent with the observed enrichment of *C. perfringens* and *Escherichia* under these conditions.

**Fig. 4.**
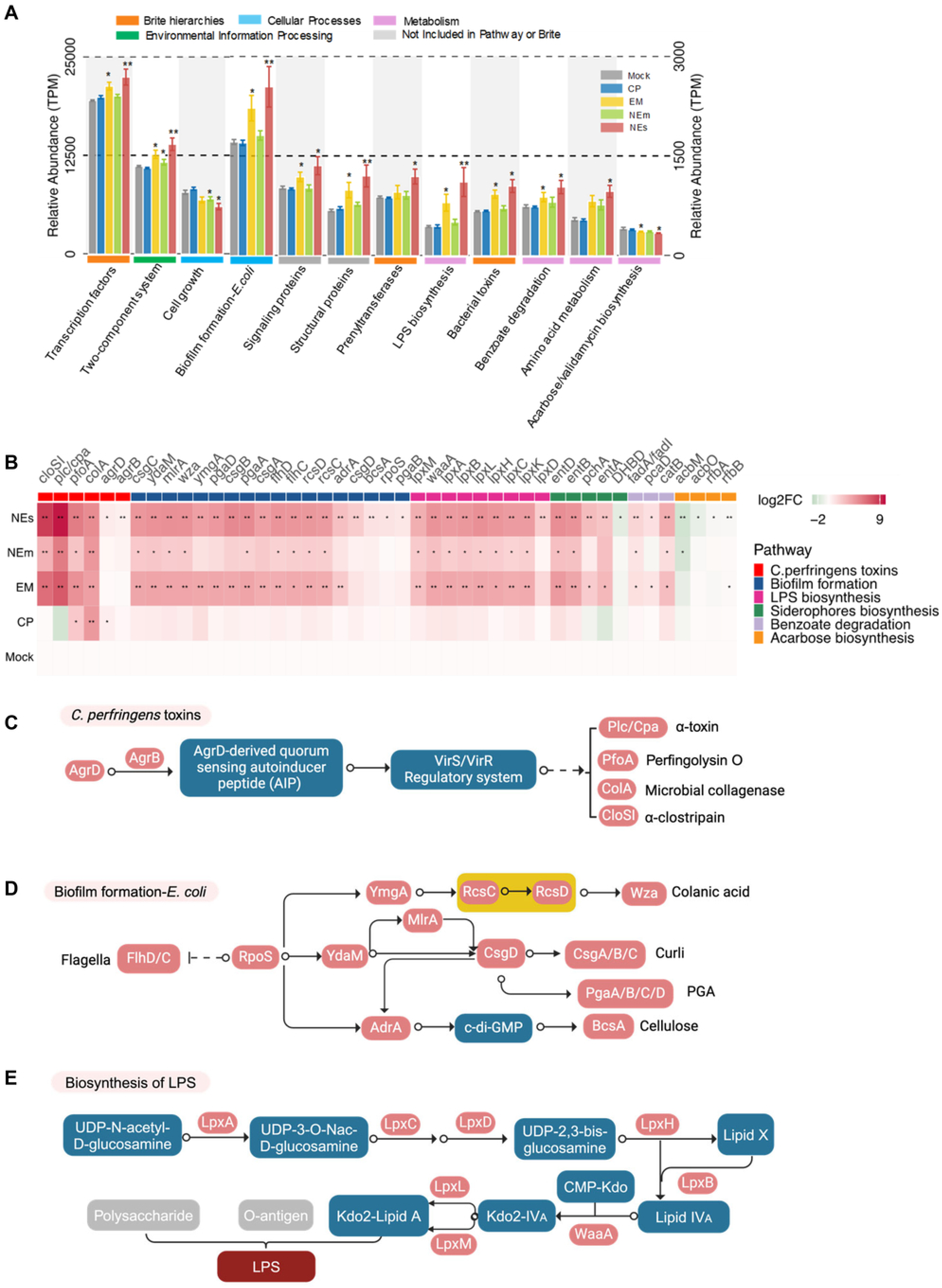
Functional alterations of the intestinal microbiota in response to infections. In Trial 1, day-of-hatch male Cobb broilers were mock-infected or orally challenged with *E. maxima* (EM), *C. perfringens* (CP), or both pathogens to induce necrotic enteritis (NE). Surviving birds in the NE group were further stratified into two subgroups: NE with mild, score-1 lesions (NEm) and NE with severe, score-6 lesions (NEs). Cecal digesta samples were collected from surviving animals (*n* = 6 per group) on day 18 and subjected to shotgun metagenomic sequencing. (**A**) Differentially enriched KEGG pathways across treatment groups. (**B**) Heatmap illustrating differential abundance of bacterial genes involved in selected pathways. Data are presented as log₂-transformed fold changes relative to mock-infected controls. In panels A and B, statistical significance was assessed using the Kruskal–Wallis test followed by post hoc Dunn’s test, with Benjamini–Hochberg correction for multiple comparisons. \**P* < 0.05, \*\**P* < 0.01, and \*\*\**P* < 0.001. **(C–E)** Pathway-specific analyses highlighting genes involved in *C. perfringens* toxin production (**C**), *E. coli* biofilm formation (**D**), and lipopolysaccharide (LPS) biosynthesis (**E**). Significantly enriched genes are shown in pink, while non-significant genes are shown in gray. Blue rectangles represent intermediate metabolites. Note that no down-regulated genes were identified in those pathways.

To further assess shifts in the cecal microbiota’s capacity for carbohydrate and particularly mucin degradation, we screened all non-redundant genes against the dbCAN2 database (14), resulting in the identification of differential abundance of the genes encoding five major CAZyme families (**Fig. 5A**). Overall, the genes encoding glycoside hydrolases (GH) and carbohydrate-binding modules (CBM) were significantly reduced in the NEs group, whereas the genes encoding glycosyl transferases (GT), carbohydrate esterases (CE), and polysaccharide lyases (PL) were significantly enriched. Subfamilies such as GH26, GH66, and CE10 exhibited a significant decrease in NEs chickens, while GH37, GH63, GH73, GH102, GH103, GT8, GT9, GT19, GT20, GT30, GT56, GT82, CBM15, CBM20, and CE1 significantly increased (**Fig. 5B**). Similar trends were observed in EM and NEm groups, albeit to a lesser extent. Notably, CP infection alone resulted in mostly no major changes in the abundance of CAZyme subfamilies.

**Fig. 5.**
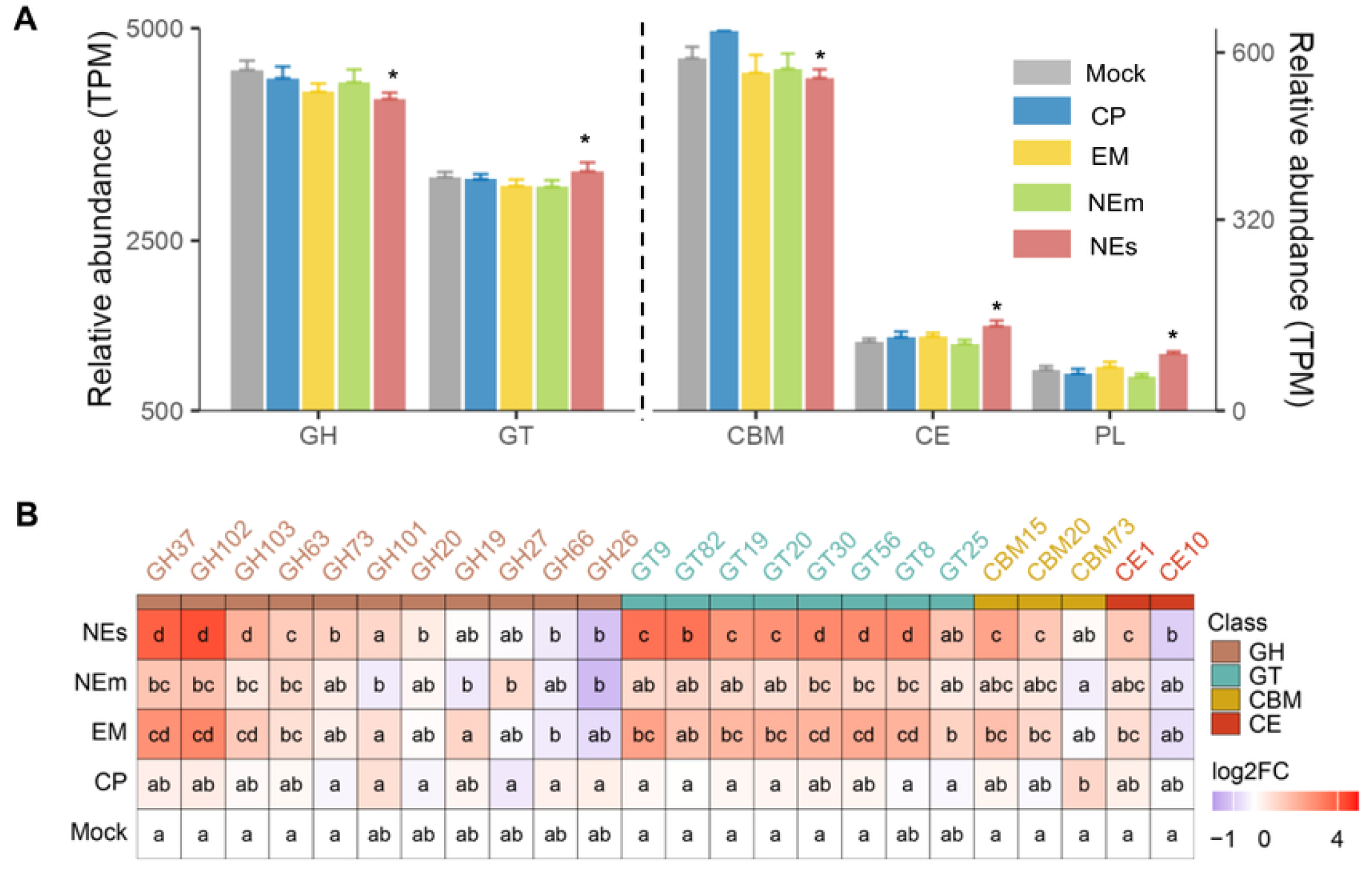
Differential enrichment of carbohydrate-active enzymes (CAZymes) in the cecal microbiota of chickens in response to infections. In Trial 1, day-of-hatch male Cobb broilers were mock-infected or orally challenged with *E. maxima* (EM), *C. perfringens* (CP), or both pathogens to induce necrotic enteritis (NE). Surviving birds in the NE group were further stratified into two subgroups: NE with mild, score-1 lesions (NEm) and NE with severe, score-6 lesions (NEs). Cecal digesta samples were collected from surviving animals (*n* = 6 per group) on day 18 and subjected to shotgun metagenomic sequencing. (**A**) Relative abundance of five major CAZyme families. Statistical significance was determined using the Kruskal–Wallis test followed by post-hoc Dunn’s test, with Benjamini–Hochberg correction. \**P* < 0.05 vs. mock-infected controls. (**B**) Differential enrichment of CAZyme subfamilies among treatment groups. Data are presented as log₂-transformed fold changes relative to mock-infected controls. Statistical significance was assessed using ANCOM-BC2 analysis. Groups not sharing a common superscript letter within a column differ significantly (*P* < 0.05).

### Temporal dynamics of the intestinal microbiota in response to CP, EM, and NE infections

To further dissect relative contributions of EM and CP on intestinal microbiota changes during the progression of NE, chickens were mock-infected or challenged with EM on day 11 and/or CP on day 15, and the ileal and cecal digesta were collected at multiple time points before and after infection (**Fig. 6A**). No discernible differences in body weight (BW) were observed among groups from days 11 to 15, but a significant reduction in BW was evident in NE-challenged chickens beginning on day 16, which persisted through day 18 (**Fig. 6B**). EM infection also resulted in significant BW loss on days 17 and 18, whereas CP infection alone had no impact on BW throughout the entire duration. The highest incidence of severe intestinal lesions occurred on day 17 in the NE group, with approximately 50% of birds receiving a severe lesion score of 6 (**Fig. 6C**). No lesions were observed in any other group throughout the study.

**Fig. 6.**
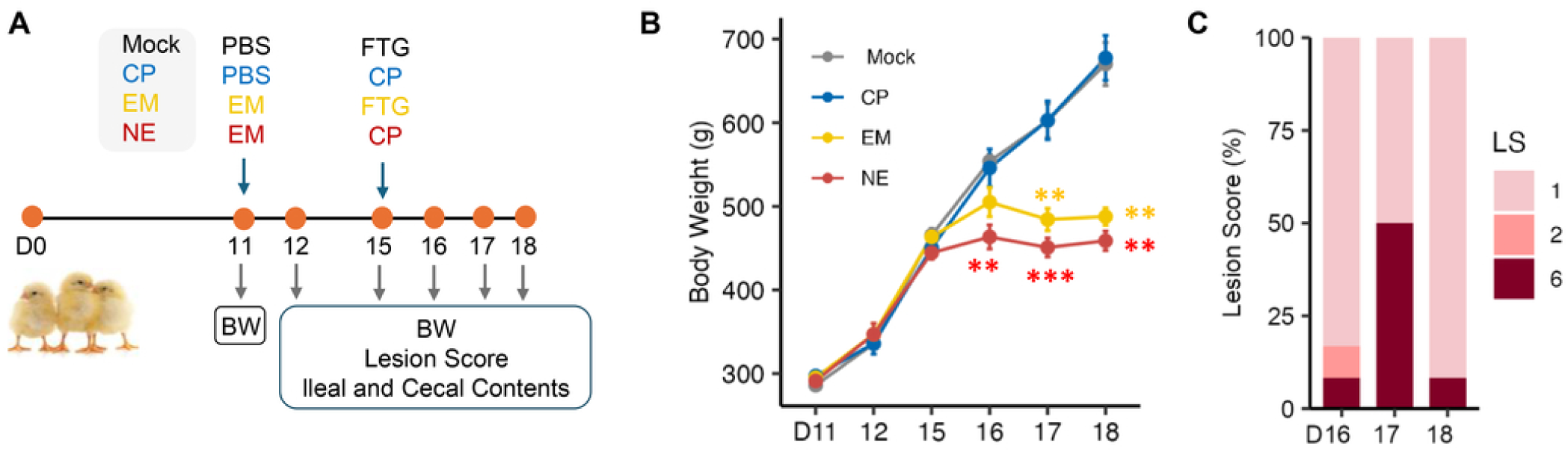
Temporal dynamics of chicken responses to infections. (**A**) Experimental design of Trial 2. A total of 200 day-of-hatch male Cobb broilers were orally challenged with *E. maxima* (EM), *C. perfringens* (CP), or both pathogens to induce necrotic enteritis (NE) on days 11 and 15, respectively. A control group received mock infections with PBS and fluid thioglycollate broth (FTG) on the corresponding days. Chickens (*n* = 10 per group) were randomly selected, weighed, and humanely euthanized for intestinal lesion scoring and collection of ileal and cecal digesta samples on various days following infection. (**B**) Changes in body weight (BW) over time following infection. Data are presented as mean ± SEM. Statistical significance was determined by one-way ANOVA followed by post-hoc Tukey test. \*\**P* < 0.01, \*\*\**P* < 0.001 vs. mock-infected birds at respective time points. (**C**) Small intestinal lesion scores (LS) in NE-affected chickens on days 16, 17 and 18. Note that no visible NE-characteristic lesions were observed in chickens from EM, CP, or mock-infected groups at any time point.

To investigate the temporal dynamics of the intestinal microbiome following infection, 16S rRNA gene sequencing was performed on bacterial DNA extracted from the ileal and cecal digesta. In the ileum, EM challenge significantly reduced Shannon Index beginning on day 15 (4 days post-infection; dpi), with effects persisting through days 16 and 17. CP infection alone caused no obvious changes at any time point, while co-infection with EM and CP resulted in a significant decrease in Shannon Index on days 17 and 18 (**Fig. 7A**). Weighted UniFrac analysis revealed significant differences in the ileal microbiota among groups from days 16 to 18 (**Fig. 7B**).

**Fig. 7.**
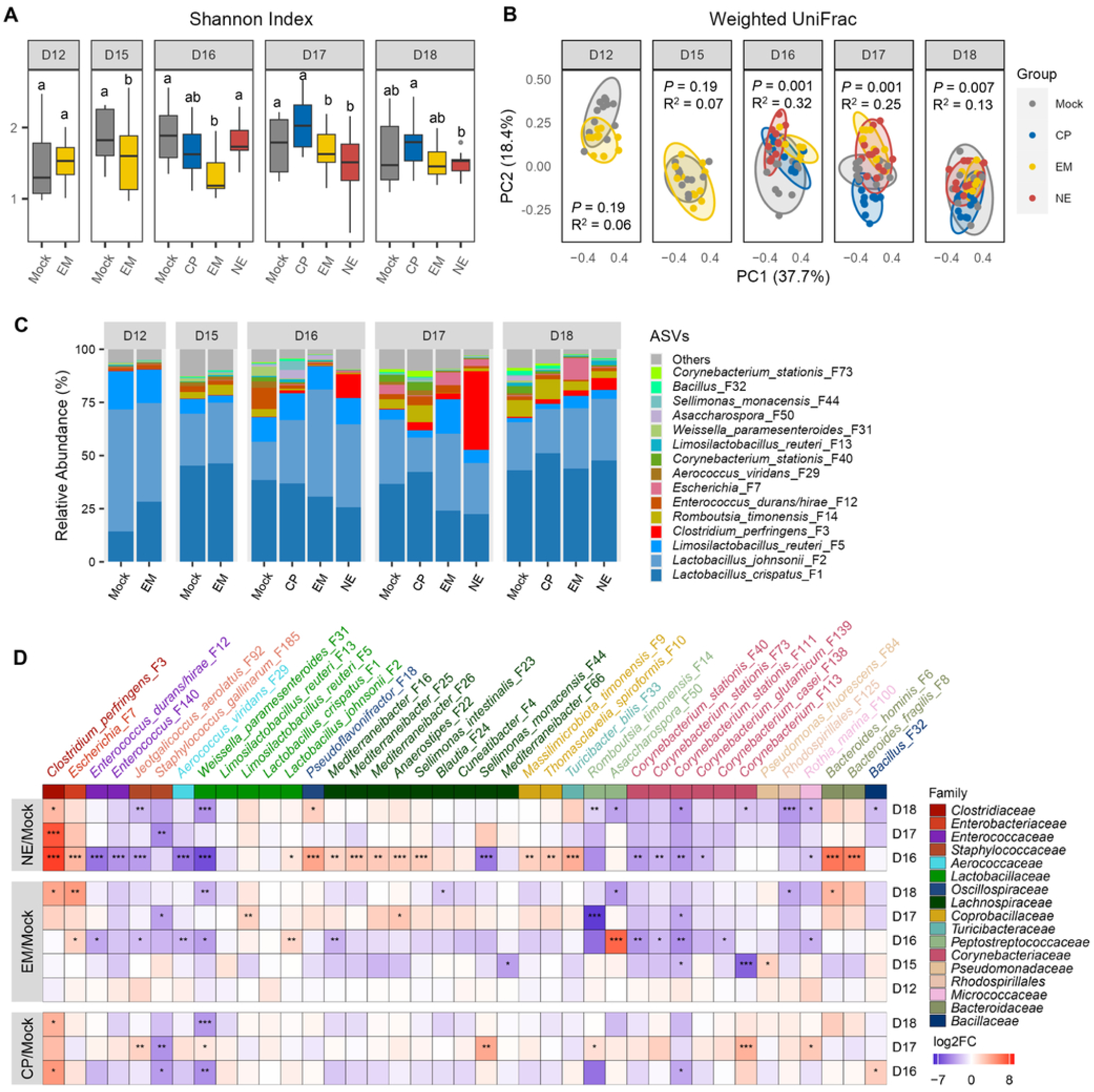
Temporal dynamics of the ileal microbiota in response to infections. In Trial 2, day-of-hatch male Cobb broilers were mock-infected or orally challenged with *E. maxima* (EM), *C. perfringens* (CP), or both to induce necrotic enteritis (NE). Ileal digesta samples were collected (*n* = 10 per group) on multiple days post-infection for DNA isolation and 16S rRNA gene sequencing. (**A**) Shannon Index visualized using box and whisker plots. Statistical significance was assessed using the Kruskal–Wallis test followed by post hoc Dunn’s test. Groups not sharing a common superscript letter differ significantly (*P* < 0.05). (**B**) Principal coordinates analysis (PCoA) plots of weighted UniFrac distances. Significance was determined using PERMANOVA with 999 permutations. (**C**) Relative abundances (%) of the top 15 bacterial amplicon sequence variants (ASVs) in the ileal microbiota. (**D**) Heatmap showing temporal changes of the top 40 ileal bacterial ASVs in response to infection. Data are presented as log₂-transformed fold changes relative to mock-infected controls at each time point. Statistical significance was determined using the Mann–Whitney U test followed by Benjamini–Hochberg correction. \**P* < 0.05, \*\**P* < 0.01, and \*\*\**P* < 0.001 vs. the mock-infected group on respective days.

Ileal microbiota composition was dynamically altered in response to infection (**Fig. 7C**). *C. perfringens* (F3) obviously increased in the NE group from days 16 to 18 and was also elevated in the CP group through the study. EM challenge led to a delayed increase in *C. perfringens* on day 18, primarily involving a netB-negative strain or a closely related *Clostridium* species, as confirmed by qPCR targeting both *C. perfringens*-specific 16S rRNA and *netB* genes (**Fig. S1A, S1B**). Distinct temporal enrichment patterns were observed among many other top ileal bacteria across groups (**Fig. 7D**). Similar to *C. perfringens*, *Escherichia* (F7) significantly increased in NE chickens on day 16, followed by a gradual decline. In EM chickens, *Escherichia* (F7) showed a progressive increase, peaking on day 18. Although *E. cecorum* (F17) was undetected, two other presumptive commensal *Enterococcus* species (F12 and F140) were significantly reduced in EM and NE groups on day 16, with F140 remaining suppressed on day 18. Among dominant LAB, *L. crispatus* (F1) slightly increased in the EM group on day 12, while *L. johnsonii* (F2) was significantly enriched in both EM and NE groups on day 16. Two *L. reuteri* strains (F5 and F13) displayed opposing trends: F5 increased in EM and NE groups starting on day 17, whereas F13 tended to decrease from day 16.

In the cecum, EM infection showed no obvious effect on Shannon Index until days 16–18. CP infection led to a transient increase on day 16, while NE had no significant impact throughout the study (**Fig. 8A**). Weighted UniFrac analysis indicated significant differences among groups beginning on day 16 **(Fig. 8B)**. Infection also caused fluctuations in the cecal microbiota composition (**Fig. 8C**). Among the top 40 cecal bacteria, *Escherichia* (F7) bloomed in CP, EM, and NE chickens starting on day 16 and remained elevated in EM and NE chickens on day 18 but subsiding in CP chickens (**Fig. 8D**). *C. perfringens* (F3) increased in CP- and NE-challenged chickens from day 16 through day 18, while EM-induced proliferation was delayed until day 17, peaking on day 18. Notably, netB-positive *C. perfringens* showed no increase in EM-infected birds, confirming the commensal origin of the bloom (**Fig. S1C, S1D**).

**Fig. 8.**
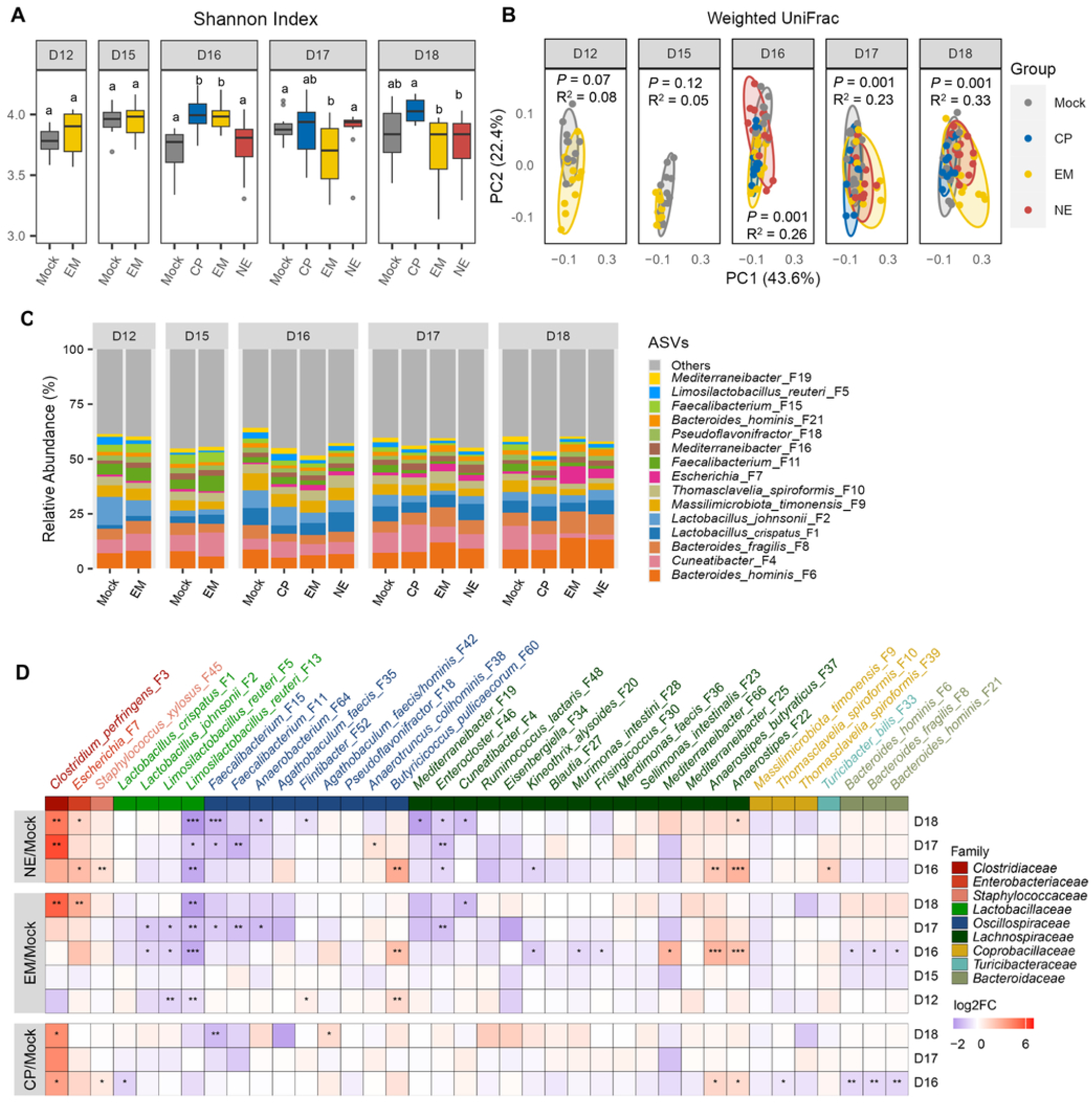
Temporal dynamics of the cecal microbiota in response to infections. In Trial 2, day-of-hatch male Cobb broilers were mock-infected or orally challenged with *E. maxima* (EM), *C. perfringens* (CP), or both to induce necrotic enteritis (NE). Cecal digesta samples were collected (*n* = 10 per group) on multiple days post-infection for DNA isolation and 16S rRNA gene sequencing. (**A**) Shannon Index visualized using box and whisker plots. Statistical significance was assessed using the Kruskal–Wallis test followed by post hoc Dunn’s test. Groups not sharing a common superscript letter differ significantly (*P* < 0.05). (**B**) Principal coordinates analysis (PCoA) plots of weighted UniFrac distances. Significance was determined using PERMANOVA with 999 permutations. (**C**) Relative abundances (%) of the top 15 bacterial amplicon sequence variants (ASVs) in the cecal microbiota. (**D**) Heatmap showing temporal changes of the top 40 cecal bacterial ASVs in response to infection. Data are presented as log₂-transformed fold changes relative to mock-infected controls at each time point. Statistical significance was determined using the Mann–Whitney U test followed by Benjamini–Hochberg correction. \**P* < 0.05, \*\**P* < 0.01, and \*\*\**P* < 0.001 vs. the mock-infected group on respective days.

Among major LAB in the cecum, *L. johnsonii* (F2) and *L. reuteri* (F5) was suppressed on day 12 and remained diminished till day 17 in EM-infected chickens, returning to baseline thereafter. CP infection appeared to enrich both bacteria on day 18, explaining a lack of significant changes in NE birds (**Fig. 8D**). *L. crispatus* (F1) was enriched early on day 12 and declined from day 16 following EM challenge, but CP infection suppressed *L. crispatus* on day 16, followed by recovery, resulting in no net changes in NE chickens. *L. reuteri* (F13) was consistently reduced throughout EM and NE challenges, with a similar but less pronounced decline in CP-infected birds. Many SCFA-producing bacteria from *Oscillospiracaceae* and *Lachnospiraceae* families such as *Faecalibacterium* (F15 and F11), *Cuneatibacter* (F4), *Mediterraneibacter* (F19), and *Enterocloster* (F46) showed progressive reductions in EM and NE chickens from day 16 onward, while CP had minimal impact (**Fig. 8D**). Conversely, transient enrichment of several SCFA producers such as *Butyricicoccus pullicaecorum* (F60) and *Anaerostipies* (F22 and F37) was observed on day 16 across all three challenge groups, but these increases largely subsided by day 18.

## DISCUSSION

NE, primarily caused by *C. perfringens*, is characterized by necrotic lesions in the small intestine of broiler chickens (2). However, *C. perfringens* alone is insufficient to induce NE, and additional predisposing factors are required to trigger clinical NE (3). Among these, *Eimeria* infection is widely recognized as the most significant contributor to NE development in commercial poultry production. *Eimeria* is a large genus of obligate intracellular parasites that invade the enterocytes upon ingestion. Approximately 3–4 days post-infection, the parasites undergo several rounds of asexual and sexual reproduction, causing rupture of epithelial cells, inflammation, and nutrient malabsorption, which peaks at 6–7 dpi following the formation and shedding of oocysts (15). Consequently, co-infection models with sequential *Eimeria* and *C. perfringens* challenges are commonly employed in experimental studies (3).

Both *Eimeria* infection and clinical NE are associated with intestinal dysbiosis, with the extent of microbial disruption closely correlating with the severity of intestinal pathology (7). Despite this, few studies have directly assessed relative contributions of *Eimeria* and *C. perfringens* to NE-associated dysbiosis. Of the four published studies addressing this question, three reported only mild intestinal lesions and no associated mortality across treatment groups (9-11), while the fourth documented a moderate mortality rate of approximately 22% in the NE-challenged group (12). These studies more closely resemble subclinical NE and may not fully capture the pathophysiological features of clinical NE, which is characterized by severe intestinal lesions, profound dysbiosis, and mortality rates that can reach up to 50% (2). Furthermore, most existing studies focused on either the ileal/jejunal or cecal microbiota, often limiting taxonomic resolution to the genus level rather than more informative ASV or species level. These limitations in low disease severity, minimum sampling sites and insufficient taxonomic resolution, have contributed to inconsistent findings across studies and hindered a clear understanding of the respective roles of *Eimeria* and *C. perfringens* in NE-induced dysbiosis and disease progression, particularly in the context of clinical NE.

In the present study, we experimentally infected chickens with *E. maxima*, *C. perfringens*, and their combination to delineate their respective impacts on the structural and functional dynamics of the intestinal microbiota in chickens with NE. Our NE co-infection model resulted in 74% lethality, with 43.6% of surviving birds exhibiting severe intestinal lesions consistent with clinical NE, and the remainder showing mild lesions indicative of subclinical NE. We analyzed structural changes of the microbiome in both the ileal and cecal samples, stratifying NE-infected birds by disease severity and enhancing taxonomic resolution down to the ASV level. Most dominant taxa were assigned to the species level using updated 16S rRNA gene databases, enabling precise identification of microbial shifts associated with both clinical and subclinical NE in the same study. Additionally, we examined functional changes of the cecal microbiota in response to CP, EM, and NE infections for the first time. The dynamics of the ileal and cecal microbiota changes were also investigated throughout the infection to study the impact of EM and CP infection on NE onset and progression.

### Relative contributions of EM and CP to dysbiosis and disease progression in NE

We observed significant changes in both the ileal and cecal microbiota in response to infection, with the ileal microbiota being more pronouncedly impacted. Notably, *E. maxima* infection, with or without *C. perfringens*, significantly reduced the richness, evenness, and overall α-diversity of the ileal microbiota, with less impact on α-diversity indices of the cecal microbiota at 7 dpi. However, EM challenge significantly affected β-diversity of both ileal and cecal microbiota. In contrast, *C. perfringens* infection alone failed to significantly alter α- or β-diversity metrics in either intestinal segment. Microbiome profiles of *E. maxima*-infected birds closely resembled those of NE-affected birds, suggesting that *E. maxima* plays a dominant role in shaping the dysbiotic landscape associated with NE. These findings are consistent with previous reports indicating that *Eimeria* infection alone or in combination with *C. perfringens* significantly reduces ileal microbiota diversity, with a less pronounced effect on the cecal microbiota (7, 9-12, 16-18). This aligns with the anatomical distribution of NE lesions, which predominantly affect the small intestine.

Temporal analysis revealed that EM infection significantly reduced α-diversity of the ileal microbiota as early as 3 dpi, with effects persisting through 7 dpi. This pattern mirrors our earlier findings on the dynamics of EM-induced dysbiosis, which typically resolves following recovery (16, 17). NE-induced dysbiosis closely resembled that observed during EM infection, while *C. perfringens* alone had minimal impact. However, because day 17 sampling included a disproportionate number of clinically symptomatic birds, the magnitude of NE-associated dysbiosis may be overestimated on day 17 and underestimated on day 18. Although EM and NE exerted less pronounced effects on the cecal microbiota, several dominant taxa in both the ileum and cecum exhibited significant and often similar alterations. Collectively, these results reinforce the conclusion that *E. maxima* is the primary driver of dysbiosis in NE, with *C. perfringens* playing a comparatively minor role.

However, NE should not be simply characterized as the result of *Eimeria*-driven intestinal microbiota disruption, with *C. perfringens* acting as a secondary factor. In our NE model, sequential challenge with both *Eimeria* and C. perfringens, rather than either pathogen alone, was required to produce the characteristic NE-associated microbial and pathological changes. Specifically, while *E. maxima* alone induced moderate intestinal pathology and a 7% mortality rate, *C. perfringens* infection alone did not result in mortality or notable intestinal lesions. In contrast, co-infection led to a dramatic increase in mortality (74%), with 43.6% of surviving birds exhibiting severe necrotic lesions (score 6) in the small intestine. These results highlight a synergistic interaction between *E. maxima* and *C. perfringens*, wherein *E. maxima* establishes a permissive environment that facilitates *C. perfringens* colonization and virulence.

The underlying mechanisms of this synergy likely involve epithelial disruption and immune modulation. *E. maxima* preferentially replicates in the jejunal and ileal epithelium, causing mucosal damage, increased mucus secretion, and the release of protein-rich nutrients that support *C. perfringens* proliferation (5). Damage to the epithelial barrier exposes extracellular matrix components, such as collagen, which serve as binding sites for *C. perfringens* adherence and toxin delivery. Additionally, *Eimeria*-induced T cell-mediated inflammation enhances mucogenesis and releases essential amino acids, further promoting *C. perfringens* growth and pathogenesis (5).

### Differential enrichment of pathobionts in response to EM, CP, and NE infections

In this study, both EM and NE challenges resulted in drastic enrichment of pathobionts, including *Escherichia*, *E. cecorum*, and *C. perfringens*, at 7 dpi. The expansion of these facultative or aerotolerant anaerobic bacteria is well-documented in inflammatory enteric diseases, such as coccidiosis (19), NE (20), and inflammatory bowel diseases (IBD) (21). Although many of these pathobiont strains are commensal and harmless in healthy hosts, under inflammatory conditions they undergo substantial expansion and may translocate across compromised intestinal barriers into tissues or circulation, thereby potentially contributing to disease processes (21).

Inflammatory reshaping of the intestinal environment appears to favor the metabolism and persistence of these pathobionts, giving them a competitive advantage over strictly anaerobic, SCFA-producing commensals (21). Co-infection with EM and CP led to a synergistic overgrowth of these bacteria, although a moderate increase was also observed in CP-infected birds. When administered independently, *C. perfringens* persisted in both the ileum and cecum at approximately 10³ CFU/g until 4 dpi, coinciding with the peak of NE symptoms. In contrast, NE-challenged birds exhibited a marked increase in *C. perfringens* load, reaching up to 10⁶ CFU/g.

To our surprise, EM-challenged birds with no prior exposure to *C. perfringens* also harbored commensal *C. perfringens* or closely related species approaching similar levels (10⁶ CFU/g). Temporal analysis revealed that while virulent *C. perfringens* persists following oral administration, commensal populations begin expanding only after 6 dpi in response to EM infection, peaking at 7 dpi. These findings suggest that EM infection promotes an intestinal environment conducive to the proliferation of *C. perfringens*, supporting its role as a key predisposing factor in NE development in commercial poultry production. Although previous studies have reported similar proliferation of *C. perfringens* and closely related *Clostridium* species in EM-challenged chickens without direct exposure to *C. perfringens* (11, 18, 22), such enrichment is not consistently observed (12, 16), likely due to inconsistent presence of *C. perfringens* in the environment or the resident microbiota.

Similar to *Escherichia spp*., *Staphylococcus xylosus* was enriched in the cecum of chickens following CP, EM, and NE challenges. *S. xylosus* is a Gram-positive, facultative anaerobe commonly found on animal skin and mucosal surfaces (23). While generally considered non-pathogenic, it has been implicated as an opportunistic pathogen in mastitis, urinary tract and septicemia (24). Interestingly, several commensal bacteria—including a *Clostridium* species (F99), *Enterococcus* (F110 and F140), and *Staphylococcus gallinarum* (F185)—were suppressed in response to infection. These taxa, belonging to the same genera as the pathobionts, may possess beneficial properties and warrant further investigation for their potential in competitive exclusion strategies aimed at mitigating NE and coccidiosis. While the precise role of these pathobionts in exacerbating clinical manifestations of NE and coccidiosis remains to be elucidated, targeting pathobionts has shown promise in IBD therapy (25), suggesting a possible translational avenue for poultry health management.

### Differential responses of LAB and SCFA-producing bacteria to EM, CP, and NE infections

The upper GI tract of chickens is predominantly colonized by LAB including major genera such as *Lactobacillus*, *Limosilactobacillus*, and *Ligilactobacillus*, as well as minor genera like *Weissella*, *Enterococcus*, and *Aerococcus* (25). In this study, we observed distinct and dynamic responses of these LAB populations to infections. Among the major LAB species, *L. crispatus* and *L. johnsonii* remained relatively stable following CP, EM, and NE challenges, although both tended to increase in EM-challenged and mildly NE-infected birds but decline in those with severe NE pathology. Interestingly, two strains of *L. reuteri* displayed opposing trends: strain F5 was consistently enriched, whereas strain F13 was diminished in both the ileum and cecum, suggesting strain-specific responses to infection, which have not been revealed in earlier studies (9-12). Minor LAB species also showed significant shifts in response to infection. *Weissella paramesenteroides* and *Aerococcus viridans*, which were undetectable in the cecum, were significantly reduced in the ileum of EM- and NE-infected birds.

Temporal analysis revealed that *L. crispatus* and *L. johnsonii* tended to increase in the ileum but decrease in the cecum at 1 dpi following CP infection and at 5 dpi after EM challenge. In contrast, *L. reuteri* (F5) shows delayed enrichment, with a significant increase in the ileum at 6 dpi following EM infection, while remaining largely unaffected by *C. perfringens*. Conversely, *W. paramesenteroides* and *A. viridans* were drastically suppressed by EM at 5 dpi and CP at 1 dpi in the ileum. The divergent responses among LAB species and particularly between two *L. reuteri* strains highlight the complexity and specificity of commensal LAB dynamics during infection. While many *Lactobacillus* and *Limosilactobacillus* species have demonstrated protective effects against NE (26), the roles of *Weissella* and *Aerococcus* in NE mitigation warrant further investigation.

The lower GI tract of chickens is dominated by various SCFA-producing bacteria capable of fermenting complex carbohydrates and modulating host immune and barrier functions (27). Among these, *Faecalibacterium* plays a pivotal role in maintaining intestinal health through its anti-inflammatory properties, enhancement of epithelial barrier integrity, and production of butyrate (28). Notably, *Faecalibacterium prausnitzii* declined significantly in the cecum at 7 dpi in both EM- and NE-infected birds, consistent with observations in coccidiosis, NE, and human conditions such as inflammatory bowel diseases and *Clostridium difficile* infection (29, 30). In addition to *Faecalibacterium*, other prominent SCFA producers such as *Cuneatibacter* and *Eisenbergiella* experienced sustained reductions from day 16 onward. However, the response of SCFA producers to infection was not uniform. For instance, *Pseudoflavonifractor* and *Anaerostipes* were markedly enriched on day 16, suggesting a potential compensatory mechanism by the host microbiota to counteract the loss of SCFA-producing bacteria. The differential dynamics of these SCFA producers in response to enteric infection remain poorly understood and warrant further investigation to elucidate their functional roles and interactions with host immunity.

### Functional alterations of the intestinal microbiota in response to EM, CP, and NE infections

Functional changes in the intestinal microbiota during necrotic enteritis (NE) remain insufficiently characterized, with most existing insights derived from predictive functional profiling based on 16S rRNA sequencing using tools such as PICRUSt and PICRUSt2 (7, 10, 31). In this study, we employed metagenomic sequencing to comprehensively characterize the functional landscape of the cecal microbiota in response to *Eimeria* (EM), *Clostridium perfringens* (CP), or combined NE challenge. Consistent with a previous functional prediction showing that NE challenge significantly increased the biosynthesis of multiple amino acids (31), we observed a significant enrichment of genes involved in amino acid metabolism in severely infected, but not moderately infected, chickens. *Eimeria* Infection likely creates a protein-rich intestinal environment through epithelial damage and impaired nutrient absorption, increasing the availability of peptides and amino acids that support bacterial fermentation and proliferation of *C. perfringens* (2, 3).

The *C. perfringens* strain used was a *netB*-positive type G isolate, selected for its well-documented involvement in NE pathogenesis via the production of key toxins, including α-toxin (Plc/Cpa), perfringolysin O (PfoA), and the pore-forming NetB toxin (32, 33). Upon reaching a threshold density, *C. perfringens* activates an Agr-like quorum sensing (QS) system, wherein autoinducing peptides (AIPs) are detected by the VirR/VirS two-component regulatory system, leading to the coordinated expression of virulence factors (34). In our data, QS-related genes, particularly *agrD* and *agrB*, were significantly enriched in birds with severe NE, suggesting robust activation of QS-mediated virulence pathways. These findings highlight the potential of QS inhibitors as therapeutic agents to mitigate NE severity by disrupting microbial communication and toxin regulation (35). Both *Plc/Cpa* and *PfoA* genes were significantly enriched not only in birds with severe NE but also in those infected with EM alone. These toxins are known to facilitate bacterial adherence and biofilm formation on exposed submucosal surfaces (32, 33).

In addition to *C. perfringens*-associated alterations, biofilm formation pathways attributed to *Escherichia coli* were markedly enriched in both EM- and NE-challenged birds. *E. coli* biofilms consist of bacterial aggregates encased in extracellular polymeric substances (EPS), which enhance survival and virulence (36). Genes encoding curli fimbriae (*csgABC*), which mediate adhesion to extracellular matrix components and abiotic surfaces (36), were significantly upregulated. Furthermore, genes involved in the biosynthesis of major EPS components—including *pgaABCD* (poly-β-1,6-N-acetyl-D-glucosamine), *bcsA* (cellulose), and *wza* (colanic acid)—were elevated in birds with severe NE. Additionally, the increased abundance of genes associated with lipid A biosynthesis, a major component of lipopolysaccharide (LPS), in EM-and NE-infected groups suggests heightened LPS-mediated host interaction, potentially contributing to immune modulation and inflammation.

Metagenomic profiling also revealed marked shifts in the abundance of genes encoding carbohydrate-active enzymes (CAZymes), particularly glycoside hydrolases (GHs), glycosyltransferases (GTs), and carbohydrate-binding modules (CBMs). GHs constituted the most abundant CAZyme class across all treatment groups, underscoring their central role in microbial carbohydrate metabolism (37). However, birds with severe NE exhibited a general decline in the abundance of GH-encoding genes, implying compromised microbial carbohydrate-processing potential. Subfamily-level analysis revealed significant reductions in the expression levels of the genes encoding GH26 (hemicellulose degradation), CE10 (xylan and pectin deacetylation), and GH66 (dextran degradation), enzymes commonly produced by fiber-fermenting taxa such as *Bacteroides* (38). These reductions point to a depletion of commensal fiber-degrading bacteria and a subsequent loss of saccharolytic functionality.

In contrast, several other CAZyme-encoding genes were significantly enriched in EM-and NE-infected chickens, including GH37 (trehalose hydrolysis), GH63 (α-glucosidase activity), GH73, GH102, and GH103 (peptidoglycan hydrolysis), GT19 and GT20 (lipopolysaccharide and exopolysaccharide biosynthesis), CE1 (xylan deacetylation), and CBM15/CBM20 (starch and xylan binding). These enzymes are predominantly associated with facultative or opportunistic pathobionts such as *Escherichia*, *Enterococcus*, *Clostridium*, and *Lactobacillus* (39). Their upregulation suggests a functional transition in the microbiota toward a community better adapted to utilize host-derived glycans and simple carbohydrates, rather than dietary fibers. This metabolic reprogramming is consistent with EM-induced mucosal damage, which increases the availability of glucose and mucus-derived substrates while reducing fiber fermentation due to impaired nutrient absorption. Collectively, these findings reflect a dysbiotic microbial state characterized by reduced complexity of carbohydrate metabolism, enhanced mucin degradation, and microbial adaptation to an inflammatory and nutrient-altered intestinal environment.

### Strengths and limitations of this study

This study has several notable strengths. The use of controlled experimental infections with *E. maxima*, *C. perfringens*, and their combination enabled clear dissection of the relative contributions of each pathogen to microbiota disruption during NE. Additionally, the NE model produced both mild (score 1) and severe (score 6) intestinal lesions, reflecting subclinical and clinical forms commonly observed under field conditions. Simultaneous characterization of compositional and functional microbiome changes in birds with mild and severe disease provided valuable insight into the pathogenesis of both NE forms. Furthermore, temporal sampling combined with high-resolution shotgun metagenomic sequencing provided detailed taxonomic and functional insights into microbiota dynamics during disease progression.

Several limitations should also be noted. First, although temporal sampling captured microbiota dynamics during disease progression, the observation window ended during peak pathology, precluding assessment of microbiota recovery following infection. Extending sampling into the recovery phase would provide valuable insight into the resilience and restoration of intestinal microbiota. While coccidiosis-induced dysbiosis appears to resolve after recovery (16, 17), it remains unclear whether the gut microbiota fully recovers following NE. Second, although shotgun metagenomic sequencing enabled detailed functional characterization, it was performed only on cecal samples. Consequently, functional alterations in the ileal microbiota, the primary site of NE pathology, were not directly assessed. Future studies integrating metagenomic or metatranscriptomic analyses of both intestinal compartments, together with metabolomic profiling, would help link microbial functional potential with metabolic activity. Third, the study employed an experimental model using *Eimeria* as the predisposing factor, which may not fully capture the complexity of field-associated NE involving additional factors such as diet, environment, and host immune status (2, 3). It would be more comprehensive if other predisposing factors were incorporated in parallel in this study. In addition, although enrichment of commensal netB-negative *C. perfringens* or closely related species was observed following *Eimeria* infection, their strain identity and functional roles remain to be determined. Addressing these limitations will help refine our mechanistic understanding of NE-associated dysbiosis and support the development of microbiome-based interventions.

## CONCLUSION

This study demonstrates that *C. perfringens* and *Eimeria* infections act synergistically to drive the onset of clinical NE, with *E. maxima* serving as the primary driver. The epithelial disruption, inflammation, and nutrient malabsorption induced by *E. maxima* replication create a permissive intestinal environment that promotes *C. perfringens* colonization, proliferation, and toxin production. Additionally, this altered niche supports the expansion of facultative pathobionts such as *E. coli*, *E. cecorum*, and *S. xylosus*, which thrive under inflammation-associated oxidative stress and exploit host-derived mucins and simple dietary carbohydrates. While dominant LAB species remain relatively stable, several beneficial, strictly anaerobic, fiber-fermenting, SCFA-producing bacteria such as *Faecalibacterium* are significantly depleted, particularly at 6–7 dpi. These microbiota shifts, including the enrichment of pathobionts and depletion of SCFA producers, offer promising targets for early NE diagnosis and therapeutic intervention. Collectively, our findings provide mechanistic insights into NE disease progression and underscore the pivotal role of *E. maxima* in facilitating *C. perfringens*-driven disease. This work lays a foundation for the development of microbiota-based diagnostic tools and therapeutic strategies for NE and potentially other enteric disorders.

## MATERIALS AND METHODS

### Chicken NE trials

Two NE trials were conducted to investigate the alternation of intestinal microbiota in response to NE using a co-infection model of *E. maxima* and *C. perfringens* as previously described (7, 40-42). All animal procedures were approved by the Institutional Animal Care and Use Committee of Oklahoma State University under the protocol number AG-21-62. In the first trial, we investigated relative contributions of *E. maxima* and *C. perfringens* to NE-induced dysbiosis. Briefly, a total of 195 unvaccinated day-of-hatch male Cobb 500 by-product breeder chicks were obtained from a Cobb-Vantress Hatchery (Siloam Springs, AR) and randomly distributed to 13 floor pens with 15 birds/pen in one environmentally controlled room. All pens were covered with fresh wood shavings, and animals had free access to tap water and an antibiotic-free standard corn-soybean meal mash starter diet (21% crude protein) throughout the study. One pen of 15 chickens was randomly assigned to the mock, *C. perfringens* (CP), or *E. maxima* (EM) group, while 150 chickens in the remaining ten pens were assigned to the NE group. After overnight fasting, chickens in the EM and NE groups were orally inoculated with 10,000 sporulated oocysts of *E. maxima* strain M6 (kindly provided by Dr. John R. Barta, University of Guelph, Canada) in 1 mL saline on day 11, while birds in the mock and CP groups received 1 mL of sterile PBS. On day 15, approximately 5 × 10^8^ CFU of *C. perfringens* strain Brenda B, carrying *netB* and *tpeL* toxin genes (kindly provided by Dr. Lisa Bielke at North Carolina State University, USA) was orally inoculated to chickens in CP and NE groups in 2 mL fluid thioglycollate broth (FTG, Thermo Fisher Scientific, Waltham, MA) twice, once in the morning and once in the afternoon with 8 h in between, while chickens in the mock and EM groups received 2 ml FTG each time.

Birds were monitored four times daily from day 11 to day 18 for behavior, clinical signs, and mortality. Chickens that were unable to stand, move, eat, or drink were humanely euthanized by CO_2_ asphyxiation to minimize undue pain. Individual body weights were recorded on days 11 and 18. All surviving animals were euthanized by CO_2_ asphyxiation on day 18, and small intestinal lesions were scored on a 6-point scale as described (13). Additionally, approximately 0.2 g of cecal digesta and 0.5 g of proximal ileal digesta were aseptically collected from each bird by gently squeezing the contents into DNase-free microcentrifuge tubes. The digesta samples were snap-frozen in liquid nitrogen and stored at −80°C until microbial genomic DNA extraction and 16S rRNA gene sequencing.

A second animal trial was conducted to determine the dynamic changes of intestinal microbiome in response to *C. perfringens*, *E. maxima*, and both pathogens. A total of 200 unvaccinated day-of-hatch male Cobb 500 by-product breeder chicks were distributed to 20 floor pens with 10 birds/pen and assigned randomly to one of four treatment groups. As in trial 1, chickens in the EM, CP, and NE groups were sequentially challenged orally with 10,000 sporulated oocysts of *E. maxima* M6, approximately 5 × 10^8^ CFU of *C. perfringens* Brenda B, or both pathogens on days 11 and 15, respectively. A control group received mock inoculations with PBS and FTG on the corresponding days. On each of the five sampling days (days 12, 15, 16, 17, and 18), 10 birds (two birds/pen) were randomly selected from each group, weighed, and humanely euthanized via CO_2_ asphyxiation. Sampling on days 12 and 15 occurred prior to EM or CP inoculation. On day 17, five less active birds were included in euthanasia and sampling. On each sampling day, approximately 0.5 and 0.2 g of the digesta were collected aseptically from the proximal ileum and cecum, respectively, snap-frozen in liquid nitrogen, and stored at −80°C until further processing. The small intestinal lesions were also examined and scored as described (13).

### 16S rRNA gene amplicon sequencing and shotgun metagenomic sequencing

Microbial genomic DNA was extracted from ileal and cecal digesta samples using the Quick-DNA Fecal/Soil Microbe Microprep and Miniprep Kits (Zymo Research, Irvine, CA), respectively. DNA concentration and quality were assessed using a NanoDrop One Spectrophotometer (Thermo Fisher Scientific). All samples achieved OD_260_/OD_280_ and OD_230_/OD_280_ ratios ≥ 1.8, indicating high quality. Deep sequencing (PE250) of the V3−V4 region of the bacterial 16S rRNA gene was performed on an Illumina NovaSeq 6000 system using primers (341F: CCT AYG GGR BGC ASC AG and 806R: GGA CTA CNN GGG TAT CTA AT). PCR amplification and library preparation were performed by Novogene (Beijing, China) using NEBNext® Ultra™ Library Prep Kit (New England Biolabs, Ipswich, MA).

Additionally, six cecal DNA samples were selected from each treatment group in the first trial for shotgun metagenomic sequencing. These samples were representative of each group, based on their 16S rRNA gene sequencing profiles. A library of 300-400 bp genomic DNA fragments was prepared for PE150 deep sequencing on a DNBSEQ platform by Beijing Genomics Institute (Shenzhen, China).

### 16S rRNA gene sequencing data analysis

Raw sequencing reads were processed with QIIME2 v2023.7 (43). Forward and reverse reads of each sample were joined, and quality control analysis was performed. Deblur algorithm v2022.8.0 (44) was used to generate amplicon sequence variants (ASVs), which were classified using the Ribosomal Database Project 16S rRNA training set v18 and Bayesian classifier (45). Taxons with bootstrap confidence of < 80% were assigned the name of the last confidently assigned taxonomic level followed by “_unclassified”. ASVs detected in < 5% of samples were excluded from downstream analyses to minimize the influence of rare taxa potentially arising from sequencing artifacts or contamination. The top 100 ASVs, along with all differentially enriched taxa, were further classified to the species level when sequence identity was ≥97%, using the GenBank rRNA/ITS database or the EzBioCloud 16S database (v2023.08.23) (46).

Analysis and visualization of α- and β-diversities of the microbiota composition were conducted using the R ‘phyloseq’ package v1.46.0 (47). The number of ASVs, Pielou’s Evenness, and Shannon Index were calculated to indicate the richness, evenness, and overall α-diversity of each microbiota sample. The β-diversity was determined using weighted and unweighted UniFrac distances (48). Differential abundance of bacteria among treatment groups was determined using ANCOM-BC2 (49).

### Shotgun metagenomic sequencing data analysis

After shotgun sequencing, all adaptors and low-quality reads were removed using Trimmomatic v 0.38 (50). Bowtie2 v2.3.4 (51) was then used to remove the reads that mapped to the chicken genome (*Gallus gallus* bGalGal1.mat.broiler.GRCg7b). Cleaned reads of each metagenome were assembled independently using MEGAHIT v1.2.9 (52). To maximize the number of metagenome-assembled genomes (MAGs) generated, co-assembly was further performed within each treatment group using MEGAHIT (52). The quality of the contigs was checked with QUAST v5.2.0 (53). Contigs from both single-sample assemblies and co-assemblies were used for metagenomic binning independently using three packages under default parameters: MaxBin v2.2.4 (54), MetaBAT2 v2.12.1 (55), and CONCOCT v1.0.0 (56). All bins generated from different methods were integrated into a superior bin set using the Binning_refiner module of MetaWRAP v1.3.2 (57). The completeness and contamination of all bins were evaluated using CheckM v1.1.6 (58) based on the lineage_wf workflow. Only the bins with ≥80% completeness and ≤10% contamination were retained. All MAGs were then dereplicated with a 99% average nucleotide identity (ANI) cutoff using dRep v3.4.2 (59). GTDB-Tk v2.3.2 (60) was used to assign taxonomy to each MAG according to the Genome Taxonomy Database (http://gtdb.ecogenomic.org/). The abundance of MAGs in each sample was estimated using metaWRAP (57) with a “quant_bins” module. A phylogenetic tree of the MAGs was built using PhyloPhlAn v3.0.6 (61) by aligning individual proteins from the protein set in each MAG and visualized using iTol v6.8.1 (62).

Contigs from each sample were selected to predict open reading frames (ORFs) using Prodigal v2.6.3 with the parameter “-p meta” (63). The predicted ORFs with a length < 100 bp were filtered out. All retained ORFs overlapping > 90% of sequence with a > 95% identity were combined to generate a nonredundant microbial gene catalog using CD-HIT v4.6.8 (64). The nonredundant gene catalogs were subjected to functional annotation using DIAMOND based on BLASTP against the KEGG (65) and eggNOG (66) databases (parameter: --evalue 10^-5^) to assign KEGG Orthology (KOs) and clusters of orthologous groups (COGs), respectively. In addition, carbohydrate-active enzymes (CAZymes) were annotated to the dbCAN2 database with a cut-off e-value of 10^-5^ using HMMER v3.1.b2 (67). Gene abundances were evaluated by mapping cleaned reads back to the gene catalog using Salmon v0.11.1 (68) and calculated in transcripts per million (TPM).

### Quantification of ileal and cecal *C. perfringens*

The ileal and cecal *C. perfringens* loads were quantified using a standard curve-based on quantitative PCR (qPCR) method as previously described (41). Briefly, bacterial genomic DNA from pure *C. perfringens* culture or intestinal digesta samples was extracted using the Quick-DNA Fecal/Soil Microbe Microprep Kit (Zymo Research) and quantified using NanoDrop Spectrophotometer. The *C. perfringens* DNA was amplified using primers specific for *C. perfringens*-specific 16S rRNA gene (AAA GAT GGC ATC ATC ATT CAA C and TAC CGT CAT TAT CTT CCC CAA A) (69) and *netB* toxin gene (GCT GGT GCT GGA ATA AAT GC and TCG CCA TTG AGT AGT TTC CC) (70) on a CFX96™ Real-Time PCR Detection System (Bio-Rad, Hercules, CA) with an initial activation at 95°C for 30 s, followed by 40 cycles at 94°C for 5 s, 60°C for 30 s and 94°C for 5 s. The *C. perfringens* titer in each sample was calculated and expressed as log_10_ CFU/g digesta based on the standard curve developed using 10-fold serial dilutions of *C. perfringens* genomic DNA.

### Statistical analysis

Statistical significance was determined using parametric or non-parametric methods depending upon data normality following the Shapiro-Wilk test. One-way ANOVA and post-hoc Tukey test was applied to compare animal weight gain, while the Kruskal-Wallis test and post-hoc Dunn’s test were used to compare α-diversity. The survival of chickens was analyzed using the log-rank test. Significance of weighted and unweighted UniFrac distances was evaluated by permutational multivariate ANOVA (PERMANOVA) using the Adonis function of the “vegan” package v2.5.6 under the default setting with 999 permutations. *P* < 0.05 or false discovery rate (FDR) < 0.05 was considered statistically significant. Differential abundance of bacteria among different groups was determined using ANCOM-BC2 (49).

## DATA AVAILABILITY

The raw sequencing reads of this study were deposited in the NCBI Sequence Read Archive (SRA) database under BioProject PRJNA1131137.

## ACKNOWLEDGMENTS

This work was supported by the USDA National Institute of Food and Agriculture grants (2022-67016-37208 and 2024-67016-42415), the Ralph F. and Leila W. Boulware Endowment Fund, and Oklahoma Agricultural Experiment Station Project H-3268. M.W. and I.T. were supported by two separate USDA National Institute of Food and Agriculture Predoctoral Fellowship grants (2021-67034-35184 and 2024-67011-42944). The funders had no role in study design, data collection and interpretation, or the decision to submit the work for publication.

The authors would like to thank Dr. John R. Barta at the University of Guelph, Canada for kindly providing *E. maxima* strain M6. The authors are also grateful to Dr. Lisa Bielke at North Carolina State University for providing the *C. perfringens* strain Brenda B.

JL, JG, MAW, IT, DMK, and GZ conducted animal trials; JL and JG processed the samples; JL, JG, and GZ analyzed the data; JL drafted the manuscript; GZ revised the manuscript; GZ conceived and supervised the study. All authors read and approved the final manuscript.

